# Computational reconstruction of clonal hierarchies from bulk sequencing data of acute myeloid leukemia samples

**DOI:** 10.1101/2021.01.23.427897

**Authors:** Thomas Stiehl, Anna Marciniak-Czochra

**Affiliations:** Institute of Applied Mathematics & Interdisciplinary Center for Scientific Computing, Heidelberg University, Heidelberg, Germany; Institute of Applied Mathematics, Interdisciplinary Center for Scientific Computing & Bioquant Center, Heidelberg University, Heidelberg, Germany

**Keywords:** Computational algorithm, acute myeloid leukemia, clonal evolution, clonal hierarchy, clonal pedigree, phylogenetic tree, bulk sequencing, stem cell

## Abstract

Acute myeloid leukemia is an aggressive cancer of the blood forming system. The malignant cell population is composed of multiple clones that evolve over time. Clonal data reflect the mechanisms governing treatment response and relapse. Single cell sequencing provides most direct insights into the clonal composition of the leukemic cells, however it is still not routinely available in clinical practice. In this work we develop a computational algorithm that allows identifying all clonal hierarchies that are compatible with bulk variant allele frequencies measured in a patient sample. The clonal hierarchies represent descendance relations between the different clones and reveal the order in which mutations have been acquired. The proposed computational approach is tested using single cell sequencing data that allow comparing the outcome of the algorithm with the true structure of the clonal hierarchy. We investigate which problems occur during reconstruction of clonal hierarchies from bulk sequencing data. Our results suggest that in many cases only a small number of possible hierarchies fits the bulk data. This implies that bulk sequencing data can be used to obtain insights in clonal evolution.

## Introduction

Acute myeloid leukemia (AML) is an aggressive cancer of the blood forming system. It is characterized by expansion of malignant cells and impairment of healthy blood cell formation [1–3]. AML originates from a small population of malignant stem-like cells, referred to as leukemic stem cells (LSC) or leukemia initiating cells (LIC). A hallmark of AML is its poor prognosis and the high rate of relapse [1–3].

The main reason for the high risk of relapse is the clonal heterogeneity of the disease. Sequencing studies reveal that the AML cell population is composed of multiple clones. Contributions of the individual clones to the total malignant cell burden vary over time [4–8]. Due to the high number of different clones, the probability is high that a subset of clones has a low sensitivity to chemotherapy, survives treatment and initiates relapse [4–9].

The clinical course of the disease shows a significant among-patient variability which can only be partially predicted based on currently existing risk-stratifications [1, 2, 9–12]. To better understand the mechanism of relapse and to identify patients at risk, a quantitative understanding of clonal dynamics is required [4–9, 13, 14].

Next-generation sequencing studies have revealed a high number of genetic hits involved in AML pathogenesis. Genetic variability among different patients is considerable and new mutations are acquired in disease evolution [4–8]. Correlation of mutations with clinical outcome has resulted in a genetics-based risk-stratification [1, 3, 15]. However, the effect of many mutations on cell dynamics remains unclear [6, 16].

Relating genetic data to patient prognosis and malignant cell properties is challenging, since different genetic hits may enhance or inhibit each other [2, 8, 9, 15–17]. Furthermore, potentially unknown or undetected hits may impact the aberrations that are observed in clinical routine. Mathematical and computational models are important to link genetic data to functional cell properties such as proliferation and self-renewal of leukemic stem cells, both of which are of prognostic relevance [9, 11, 12, 13, 14, 18, 19].

Such models allow to estimate which leukemic cell properties correspond to the clinical course of an individual patient and to link the estimates to mutation data [9, 11, 12]. This provides insights into the impact of different mutations and leads to new hypotheses about the underlying biological mechanisms and genotype-phenotype correlation.

Leukemic stem cell dynamics are governed by two key properties: proliferation rate and fraction of self-renewal. The proliferation rate describes how often LSC divide per unit of time. Upon division a LSC gives rise to two progeny, which can either be LSC or of a more differentiated progenitor type. The fraction of self-renewal corresponds to the fraction of LSC among the progeny [20, 21]. Mathematical and computational models suggest that stem cell properties at diagnosis differ from those at relapse. Particularly, LSC at diagnosis are characterized by an increased self-renewal fraction and a higher proliferation rate compared to healthy cells. LSC at relapse are characterized by a slow proliferation rate and a further increase of the self-renewal fraction [9, 19]. Computer simulations and model analysis indicate that increased self-renewal leads to a competitive advantage of the respective clones and that clones appearing later in the course of the disease have a higher self-renewal compared to clones emerging earlier [9, 13, 14, 19, 22].

Single cell sequencing technology allows to detect mutations that are present in a single cell. Sequencing of a sufficiently large number of single cells allows to reconstruct the order of mutation acquisition and to visualize it as a so-called clonal hierarchy, clonal pedigree or phylogenetic tree [7, 23]. Computational models have led to the hypothesis that the position of a clone in the phylogenetic tree correlates with its fraction of self-renewal [19]. Therefore, phylogenetic trees may contain important information about cell properties that could be used to decipher the impact of mutations on the malignant cell kinetics.

In contrast to the single cell sequencing approach, bulk sequencing analyses a mixture of DNA of multiple cells, to which each cell contributes its specific (either mutated or non-mutated) alleles. Since in most cases each cell carries two versions of each allele, the bulk sample from *n* cells is a mixture of 2*n* allele versions. The so-called variant allele frequency (VAF) is the percentage of allele versions that is mutated. Bulk sequencing quantifies the frequency of a mutated allele in a cell population however does not determine how the detected mutations are distributed among the different clones [23, 24, 25].

Single cell sequencing is a relatively new and costly technology that so far is not used in clinical routine [25]. To deduce clinically relevant knowledge from genetic data large patient groups have to be studied due to the high inter-individual heterogeneity of the detected mutations and their unknown interaction. For this reason, it is a relevant question whether clonal hierarchies can be deduced from bulk sequencing data which are routinely obtained after initial diagnosis of AML [24, 25], although most of the diagnostic sequencing is targeted on limited panels of “typical” driver mutations.

In this work we propose an algorithm that systematically constructs *all* phylogenetic trees that are in agreement with bulk sequencing data of an individual patient. This algorithm provides a tool to better understand the ambiguity of such reconstructions and their sensitivity to measurement errors.

To test our approach, we choose a recently published set of single cell sequencing data as a gold standard (ground truth) [7]. Based on the single cell sequencing data we calculate the variant allele frequency of the different mutations in a bulk sequencing sample and test whether the ‘real’ clonal pedigree, i.e., the pedigree deduced from single cell sequencing data, can be reconstructed from it. We investigate how the correctness and uniqueness of the reconstruction depend on sampling and measurement errors.

## Materials and Methods

### Aim

We use variant allele frequencies from bulk sequencing as input data. The output we want to obtain are all clonal hierarchies that are compatible with the input data.

### Assumptions

We assume that each mutation is only acquired once. Variant alleles cannot mutate back to wild type alleles. We only consider heterozygous mutations. We rescale the measurements such that the variant allele frequency of the most abundant mutation is equal to 100%.

### Computational Methods

The method is summarized in Fig. 1. Assuming that each mutation is irreversible and only acquired once, clonal pedigrees have the structure of labelled rooted trees. An (unrooted) tree is a undirected acyclic connected graph [26]. If one node of the tree is designated as root, a rooted tree is obtained. In a rooted tree we naturally assign directions to the edges pointing from the root towards the leaves. If a unique label is assigned to each node, the tree is referred to as a labelled tree [26]. The root of the tree corresponds to a genetic trait that is present in all clones. If the disease originates from a single founding mutation that is present in all malignant cells, the root can be identified with the founding mutation. This configuration applies to most leukemic patients. If there exist multiple founding mutations the root of the tree corresponds to the healthy phenotype. Each node in the tree corresponds to one clone. The label assigned to a node indicates the mutational events that gave rise to the clone. The edge pointing towards the node indicates which ancestor clone acquired the mutational event indicated by the label.

**Figure 1:**
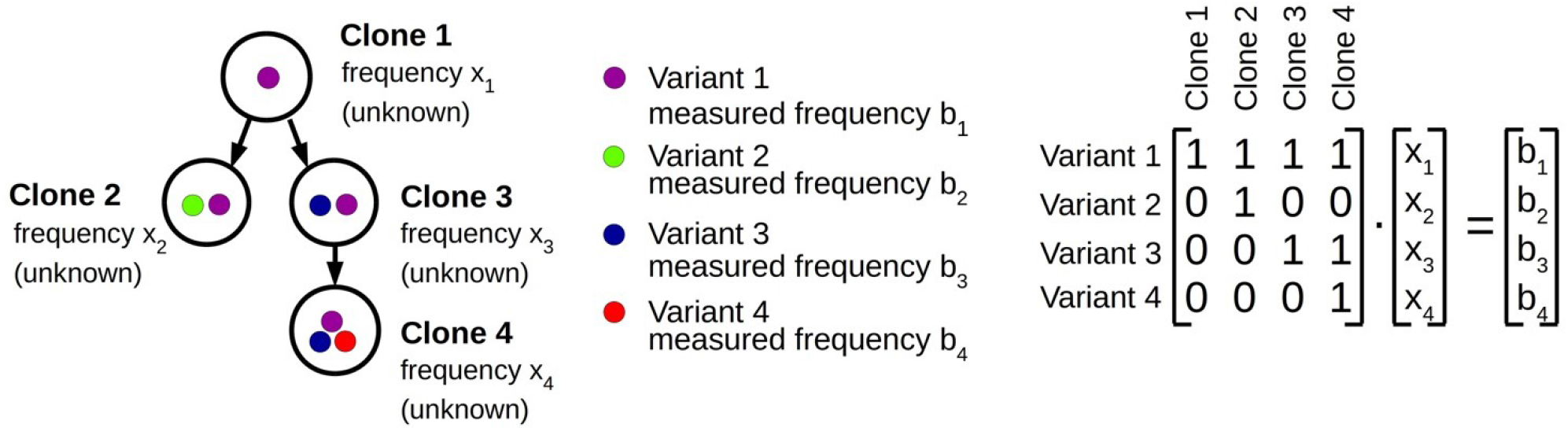
Computational approach. Clonal hierarchies are rooted trees. The root of the tree either corresponds to wild type cells or to the AML founder clone. The coloured dots represent different alleles or mutations. From bulk sequencing the allele frequencies *b_i_* in the sample are known. The frequencies *x_i_* of the different clones are unknown. The tree structure is represented by a triangular matrix. The measured data is compatible with the tree structure if the system *Ax=b* has a non-negative solution.

The tree structures can be mapped to matrices. We consider a tree with *n* nodes, corresponding to *n* clones denoted by *clone* 1 to *clone n*. Since each clone differs from its ancestor by exactly one new mutation, there exist *n* different mutations, which we number from 1 to n. Denote by *A_1≤i,j≤n_* a matrix. We set *a_ij_*=1 if clone *j* carries mutation *i*, otherwise we set *a_ij_*=0. We number the clones starting from the root (=*clone* 1) and proceed with increasing depth, i.e. if the depth of *clone i* is higher than the depth of *clone j*, then *j* We denote the founding mutation as *mutation* 1 and the mutation that is present in *clone j* but not in its direct ancestor as *mutation j.* Then *A_1≤i,j≤n_* is an upper triangular matrix, with *a_ii_*=1, and *a_ij_* from the set {0,1}.

We aim to solve the linear system of equations *Ax=b*, where *b_i_* is the measured frequency of *mutation i* in the bulk sample and *x_i_* is the abundance of *clone i* in the sample. We note that A has determinant 1 and therefore this system of equations has a unique solution. The solution is biologically feasible if all *x_i_* are non-negative. The existence of a non-negative solution can be easily checked since the solutions of *Ax=b* are given by *x_n_*=*b_n_, x_j_*=*b_j_-a_jj+_*_1_ *x_j+1_ - … - a_jn_x_n_*. We say that the dataset *b* is compatible with the clonal hierarchy represented by matrix *A* if *Ax*=*b* has a non-negative solution.

The founder mutation is denoted as *mutation* 1. It is present in all clones and, therefore, is the most abundant mutation in the bulk sample. This implies that *a_1j_=*1 for 1≤*j*≤*n.* Since we normalized the frequency of the most abundant mutation to 100% it holds *b_1_*=100. This implies that the sum over the *x_i_* is equal to 100.

To systematically generate all possible trees, we use Pruefer sequences, a classical concept to bijectively map unrooted trees with *n* nodes to sequences of length *n*-2 [27]. Each unrooted labelled tree with *n* nodes then corresponds to a sequence of length *n*-2 with elements from {1, …, *n*}. This implies that there exist n^n-2^ unrooted labelled trees. Since each of the *n* nodes can be designated as root, there exist n^n-1^ labelled rooted trees.

### Interpretation

If a biologically feasible solution of the system *Ax*=*b* exists, the measured bulk allele frequencies *b* can be explained by the tree structure that corresponds to the matrix *A*. This means that the bulk allele frequencies *b* are obtained by mixing the different clones from the tree in appropriate proportions (the abundance of *clone i* has to equal *x_i_*). For each pair *A, b* a biologically feasible solution can exist or cannot exist. For example, a tree with founder mutation *X* (i.e., each clone carries mutation *X*) cannot match to samples where the abundance of *X* in non-maximal.

### Measurement Errors

If the measured data *b* are exact, non-existence of a biologically feasible solution indicates a mismatch of the tree structure and the allele frequencies. In case of experimental data, the non-existence of a biologically feasible solution can alternatively arise from measurement errors. For this reason it may be necessary to also consider solutions fulfilling *||Ax-b||<ε* for an appropriate *ε*, where *||.||* denotes e.g., the Euclidean norm.

To find such solutions, especially in the case where no biologically feasible (i.e., exact non-negative) solutions exist, we use an optimization approach to obtain a non-negative solution that reproduces the data as good as possible. For each matrix *A* that corresponds to a tree structure we minimize ||Ax-b|| under the constraints *x_i_*≥0 (*i*=1,…n), *x_1_+…+x_n_*=100. If the measured VAF have different confidence intervals, we minimize the weighted error function *||W*(*Ax*-b)*||*, where *W* is a diagonal matrix with entries related to the confidence intervals.

Solving the minimization problem for each possible tree structure allows to rank the tree structures based on the mismatch *||Ax-b||* and to identify which tree optimally fits to the data. A solution is referred to as exact if |*|Ax-b||*<10E-16. We say that the tree structure corresponding to matrix *Ã* is optimal if it holds *||Ãx*-*b||*≤*|Ax*-*b||* for all matrices *A* that represent a suitable tree and vectors *x* fulfilling *x_i_*≥0 (*i*=1,…n), *x_1_+…+x_n_*=100. The optimization was carried out using the python cvxopt package [28].

### Data

We plan to investigate if it is possible to reconstruct clonal hierarchies from bulk sequencing samples. This requires that the “true” clonal hierarchy is known, so that we can compare the result of our algorithm with reality. To know the “true” hierarchy we use single cell sequencing data from ref. [7]. We understand the clonal hierarchy and the clonal frequencies obtained from the single cell sequencing as ground truth. Since for the samples analysed in [7] no bulk data are available, we calculate the bulk allele frequencies based on the single cell data. For simplicity we assume that the considered sample only contains leukemic cells and we exclude all sequenced wild type cells from the data. We calculate the bulk VAF of variant allele *i* as *a_i1_f_1_+…+a_in_f_n_,* where *f_i_* is the frequency of clone *i* in the single cell data set and *a_ij_*=1 if clone *j* carries variant allele *i* and 0 otherwise. Since we consider a purely leukemic sample, the calculated VAF are normalized such that the frequency of the most abundant variant allele is 100%. We consider all patients from ref [7] that carry heterozygous mutations and for whom data at diagnosis and relapse is available.

## Results

### Exact input data often result in unique clonal hierarchies

As gold standard we use the single cell sequencing data from ref. [7], which provide the true clonal hierarchy and hence can be used to test the proposed algorithm. Based on the single cell data we calculate the variant allele frequencies in the bulk sample. The first question we ask is how many clonal hierarchies are compatible with the bulk variant allele frequencies of a given patient. Fig. 2 shows for each patient which hierarchies exactly fit to the data at diagnosis. We observe that, for 5 out of 6 patients, only one hierarchy exactly fits the bulk data. For one patient 6 hierarchies are consistent with the bulk data.

**Figure 2:**
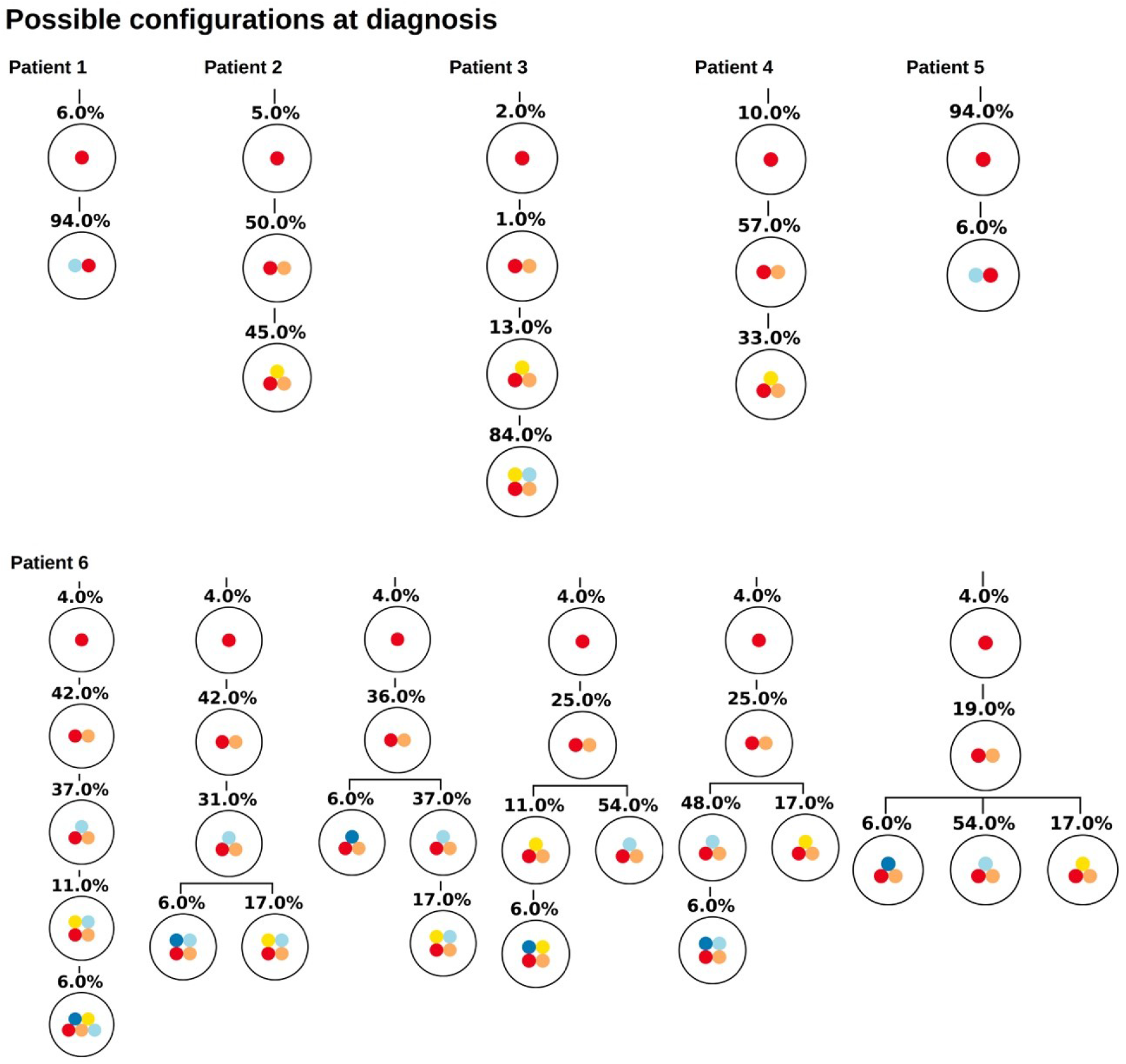
Clonal hierarchies compatible with the bulk allele frequencies measured at diagnosis. For each considered patient all clonal hierarchies are depicted that are compatible with the bulk variant allele frequencies measured at diagnosis. The root of the tree corresponds to the founder mutation that is present in all leukemic cells. The percentages indicate the frequencies of the respective clones that have to be mixed to obtain the measured bulk VAF. We observe that in most cases the hierarchies are unique.

Similar observations hold for the relapse samples of the considered patients, Fig 3. Here all samples lead to unique tree configurations. In the next step we combine the diagnosis and relapse sample of each patient. For each patient Fig. 4 shows the tree configurations that are compatible with the data at both time points. Again, we have uniqueness in all except one case.

**Figure 3:**
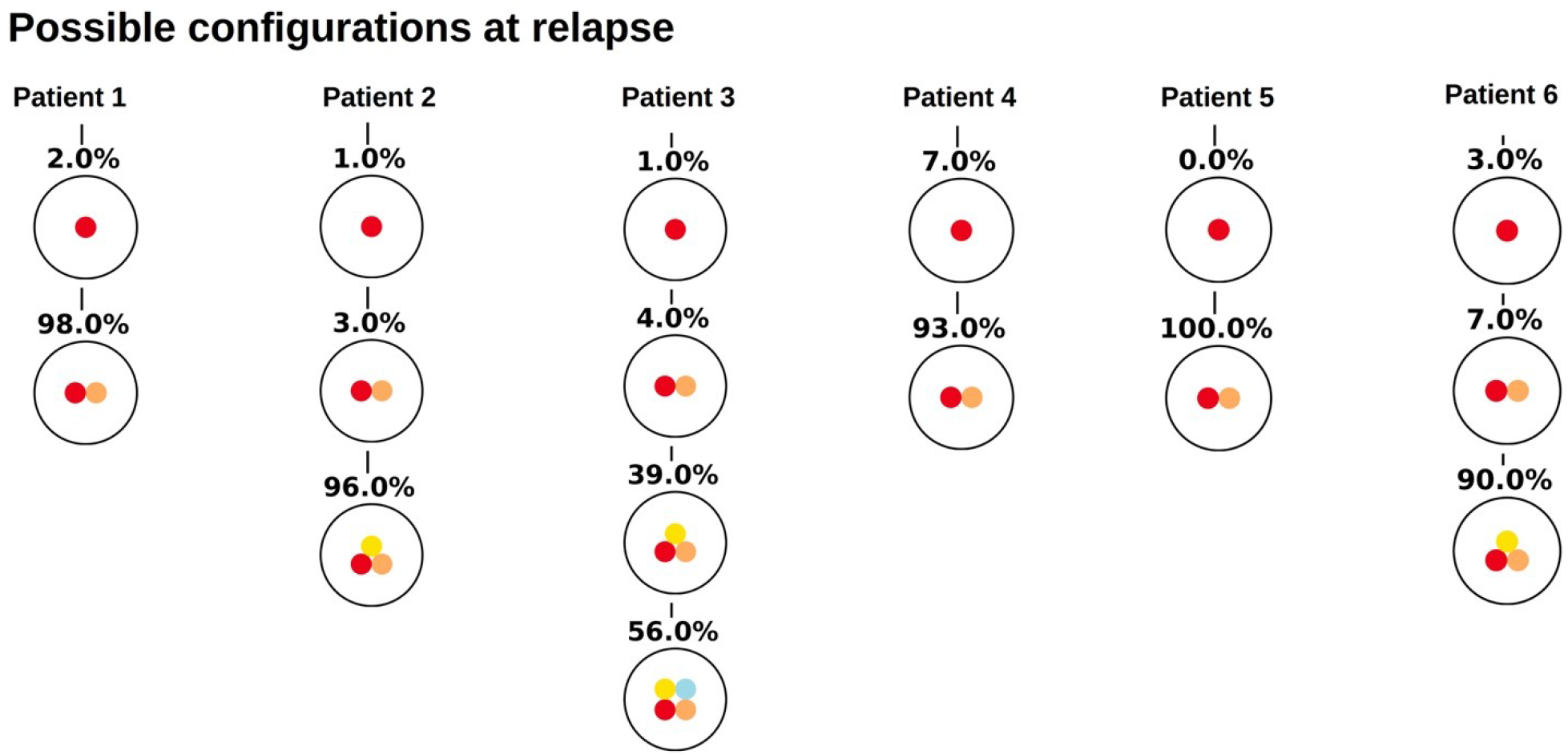
Clonal hierarchies compatible with the bulk allele frequencies measured at relapse. For each considered patient the Figure shows which clonal hierarchies are compatible with the bulk variant allele frequencies measured at relapse. The root of the tree corresponds to the founder mutation. We observe that in all cases the hierarchies are unique.

**Figure 4:**
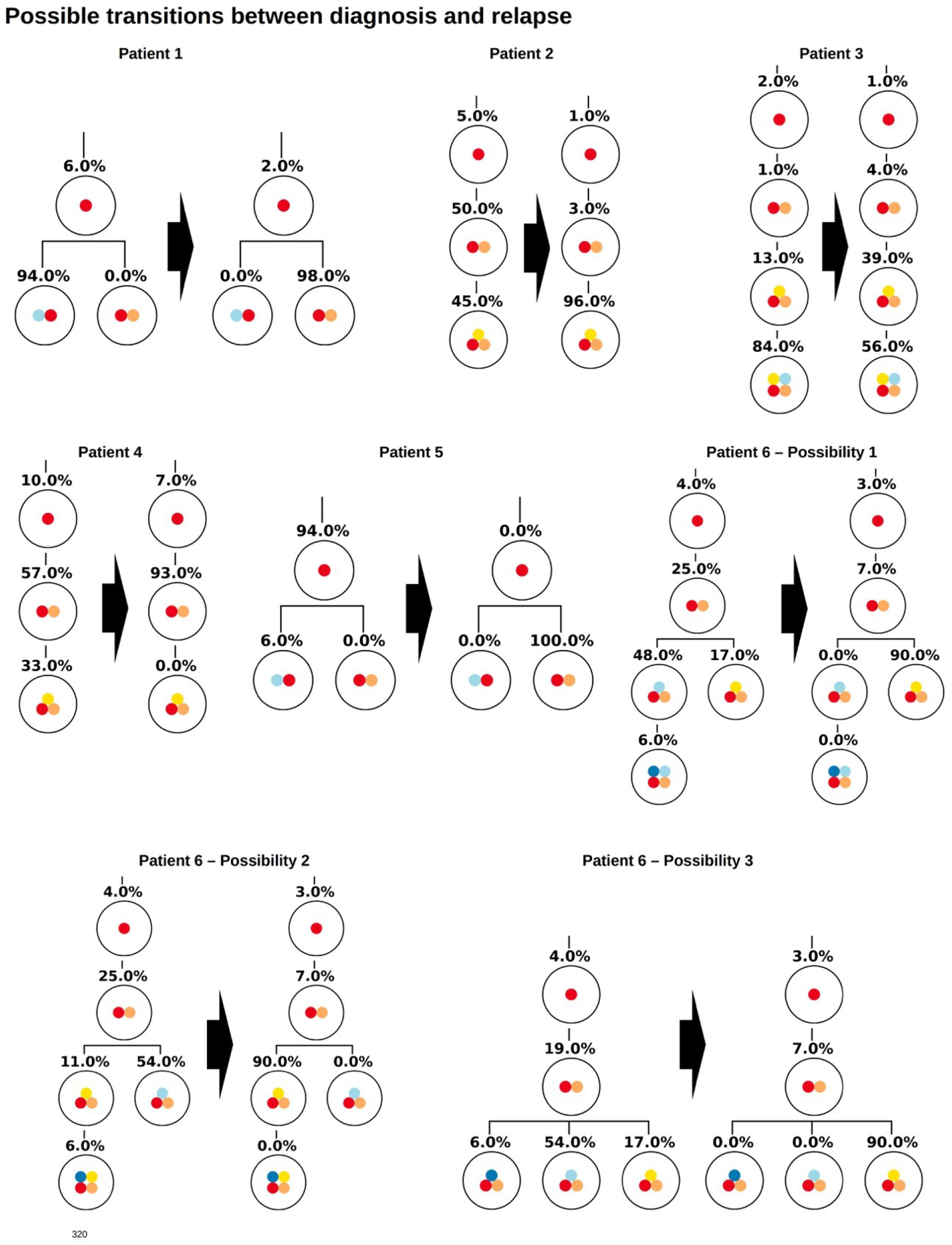
Clonal hierarchies compatible with the bulk allele frequencies measured at diagnosis and relapse. For each patient the Figure shows all clonal hierarchies that are compatible with the bulk VAFs measured at diagnosis and relapse. The root of the tree corresponds to the founder mutation. We observe that in case of patient 6 the number of hierarchies compatible with the data is reduced compared to Figure 2. For patients 1-5 the reconstructed hierarchies coincide with the result from single cell sequencing. For patient 6 Possibility 3 corresponds to the true configuration.

### Sampling error has little impact on the uniqueness of clonal hierarchies

If the frequency of different clones in a large population is estimated based on a small sample, sampling errors can occur. To study the impact of sampling errors on the reconstructed clonal hierarchies we again use the single cell sequencing data from [7]. We assume that the single cell data reflect the true frequencies of the clones in the malignant cell bulk of the respective patient. For an arbitrary patient *k* we know the total number *n_k_* of sequenced leukemic single cells. Furthermore we know the frequencies *f_i,k_* of each clone that has been detected (here *f_i,k_* denotes the frequency of clone *i* in the sample of patient *k*). To study the impact of sampling on the bulk variant allele frequencies and on the reconstructed hierarchies for patient *k,* we draw 1000 random samples of size *n_k_* from a multinomial distribution with probabilities *p_i_=f_i,k_*. This approach is referred to as resampling [29].

For each of these 1000 random samples we calculate the bulk variant allele frequencies and apply our algorithm to reconstruct the clonal hierarchies. The results are shown in Fig 5. In all cases except one the hierarchies fitting exactly to the data remain unique and are identical to the hierarchies obtained based on the exact data. For one patient in some of the resampled datasets the number of hierarchies matching the data increases by one. These results imply that the sampling error has a negligible impact on the clonal pedigrees that fit to the data.

**Figure 5:**
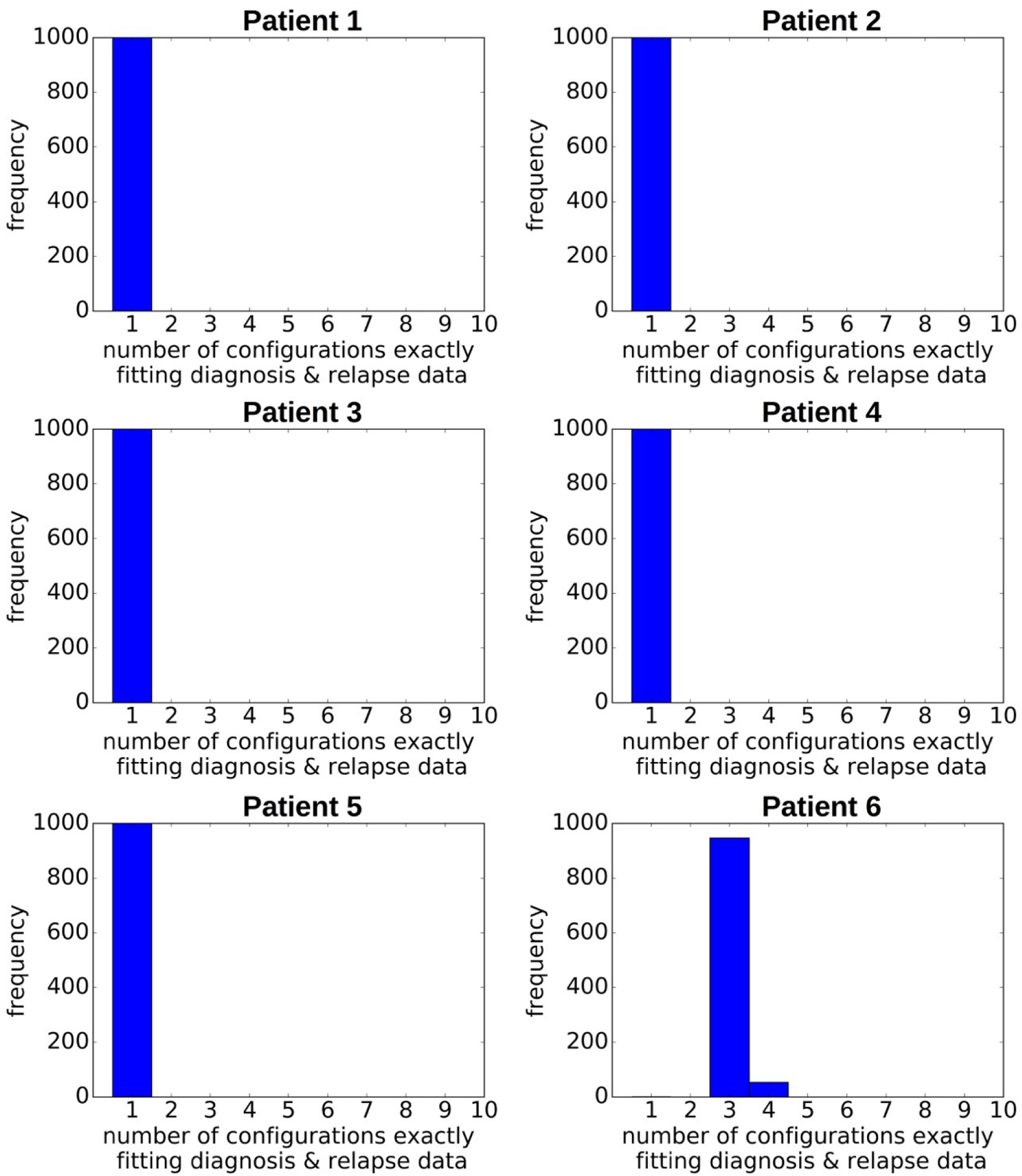
Impact of sampling error on the reconstructed hierarchies. For each patient we generate 1000 random pairs of diagnosis and relapse samples from a multivariate distribution. The probabilities of the multivariate distribution equal the clonal frequencies in the single cell data. The size of the samples equals the number of sequenced leukemic cells. We recorded for each pair of randomly generated diagnosis/relapse samples the number of clonal hierarchies compatible with the resampled diagnosis and relapse data. The vertical axis shows how many of the 1000 samples were compatible with 1, 2, 3, … hierarchies respectively.

### Impact of measurement errors on reconstruction of clonal hierarchies

Inaccuracies in sequencing are another possible source of error. To study their impact on the reconstructed clonal hierarchies, we add a normally distributed error to the bulk frequency of each allele. Such errors can have different impacts on the reconstructed clonal hierarchies. For each patient we considered 1000 randomly perturbed versions of the original data. If the standard deviation of the error distribution is 0.5% (i.e., in 68% of cases the error is less or equal 0.5%, in 95% of cases the error is less or equal to 1%) the reconstruction algorithm works reliably in the sense that the true configuration is an optimal configuration, see Fig 6. In 5 out of 6 considered patients the optimum is unique. We repeated the simulation for a normally distributed error with a standard deviation of 5%, i.e., in 95% of cases the error is less than 10%, see Fig 6. For an error of this magnitude the true configuration not always remains an optimal configuration. This especially applies to patients in whom the frequency of the founding clone is small (i.e., patients 2, 3 and 6). If the error is larger than the frequency of the founder clone it becomes impossible to reliably detect which hit occurs first. However, also in a single cell sequencing approach, rare clones can remain undetected due to sampling or sequencing errors, implying that the first hit remains unknown. In terms of variant allele frequencies this implies that trees cannot be reliably reconstructed if the difference between the two most abundant allele frequencies is of the order of magnitude of the sequencing error. In patients with many clones, our algorithm can often rule out most of the possible hierarchies and identify a small number of configurations fitting the data. In case of Patient 5 the true configuration is always among the upper 12% of the best fitting configurations (i.e., the best or second best), and in patient 6 among the upper 3.3% of the best fitting configurations (i.e., 4 out of 125). In case of small clone numbers such as for patients 2 or 4, the true configuration is always among the two best fitting hierarchies.

**Figure 6:**
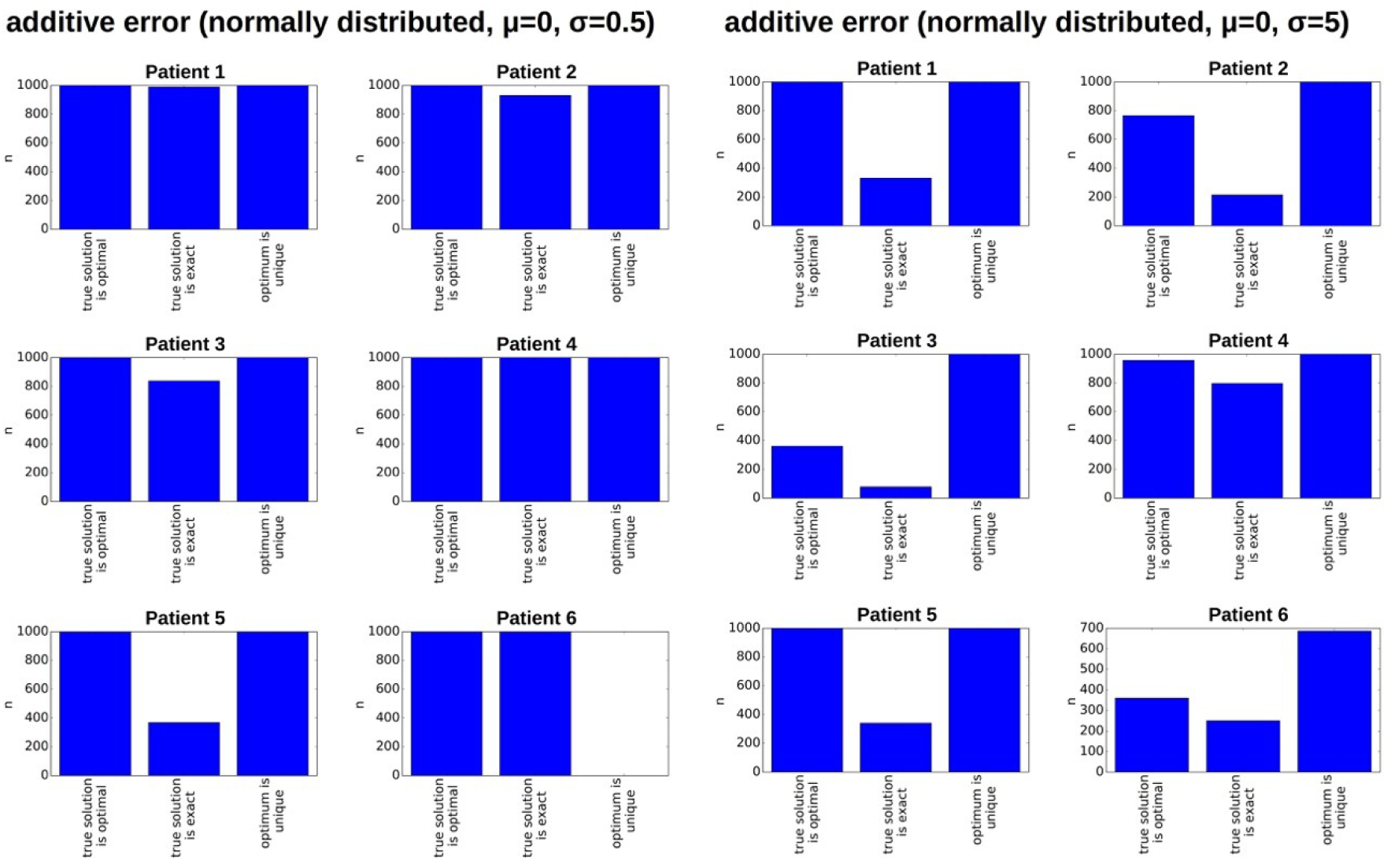
Impact of measurement errors on the reconstructed hierarchies. For each patient we generate 1000 randomly perturbed pairs of diagnosis and relapse samples. The additive random perturbations were drawn from a normal distribution with mean zero and standard deviation 0.5 or 5 respectively. Perturbations leading to a VAF of less than zero or more than 100% were excluded. For each of the perturbed diagnosis/relapse pairs we reconstructed all compatible clonal hierarchies. The Figure indicates for how many of the 1000 perturbed samples the true hierarchy optimally fits the perturbed data (compared to all other existing hierarchies), whether the true hierarchy can exactly reproduce the perturbed data and whether the optimal configuration is unique.

### An example of a patient with two founder clones

We now consider an example of a patient with two different founder clones. This scenario either corresponds to the rare case where the AML cell population originates from clones with different initial mutations or it corresponds to the case where the common founding mutation has not been detected. The latter may especially occur in the setting of targeted sequencing, where only a predefined subset of mutations is considered. Such a scenario occurs if in a purified AML sample (i.e., in a sample without healthy cells) all bulk VAF are significantly different from the expected maximum of 50% (for heterozygous mutations) or 100% (for homozygous mutations). The proposed algorithm can take this scenario into account by considering healthy cells as the root of the tree. This means a healthy reference allele that is present in all cells is added to the list of variant allele frequencies, to obtain a single tree with a unique root. Fig 7 shows all tree structures that are compatible with the measured data. The tree structures can be divided into two classes. In the first class of solutions, the frequency of healthy cells is zero at diagnosis and relapse (possibilities 1-2), in the second class the frequency of the healthy cells is positive (here 15%) at at least one time point (possibilities 3-7). Solutions of the first class imply that there exist two founding clones (or an undetected unique founder mutation), solutions of the second class may imply that the sample contains a mixture of healthy and leukemic cells. If we can be sure that the experimental procedures prevent healthy cells from being sequenced (e.g., by FACS sorting for a leukemia specific surface marker before sequencing), only two possible tree structures remain.

**Figure 7:**
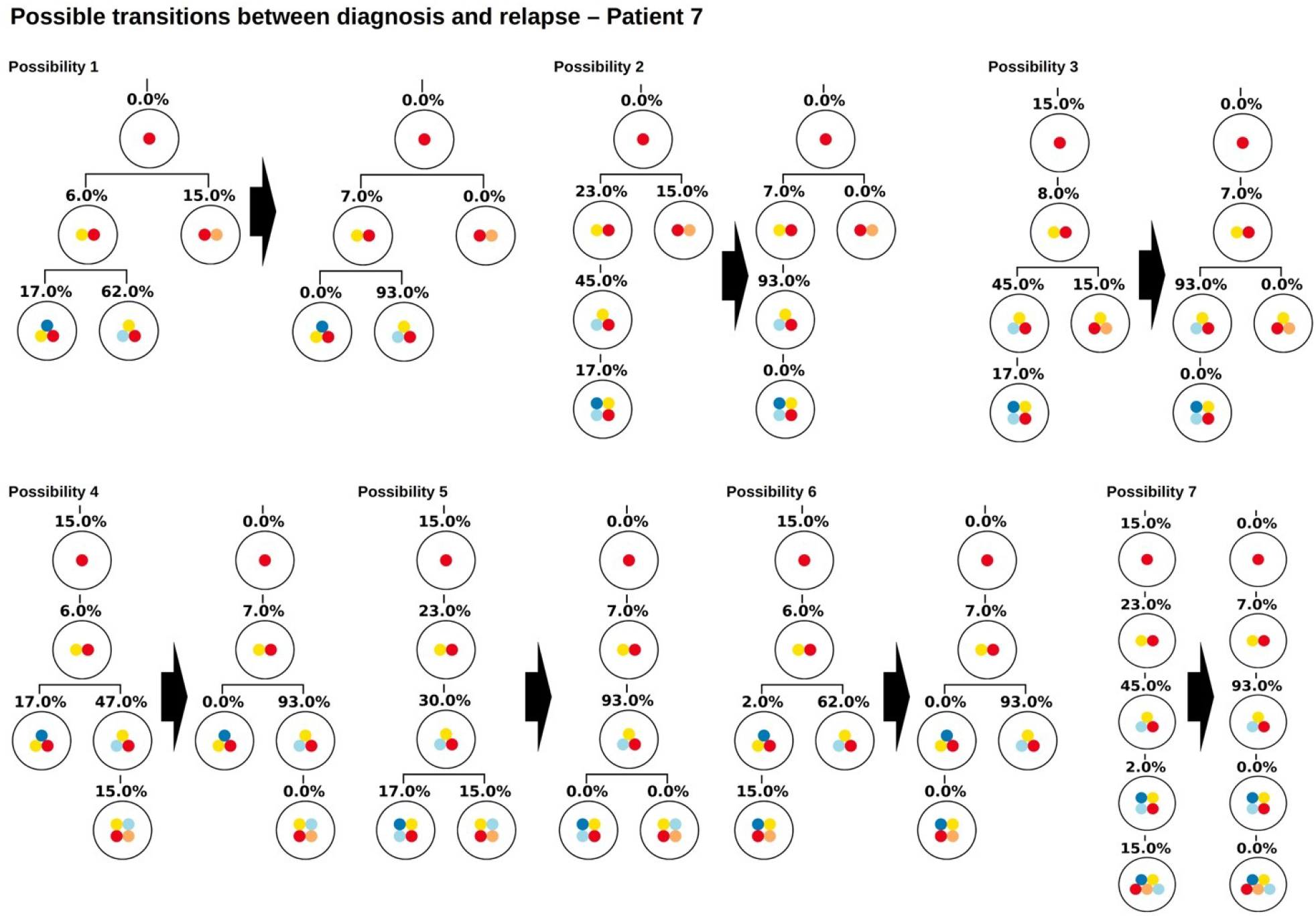
Example of a patient with multiple founder clones. In this Figure the root corresponds to wild type cells. The two founding events are indicated by yellow and orange circles. The Figure shows all hierarchies that are compatible with VAFs at diagnosis and relapse. Possibility 1 coincides with the hierarchy obtained from single cell sequencing.

As for the other patients the sampling error only leads to small changes in the numbers of clonal hierarchies that fit the data, Fig 8 (A). However, already small errors added to the bulk VAFs (normally distributed with a standard deviation of 0.5%) imply that in a majority of cases the true solution is no longer optimal, Fig 8 (B, C).

**Figure 8:**
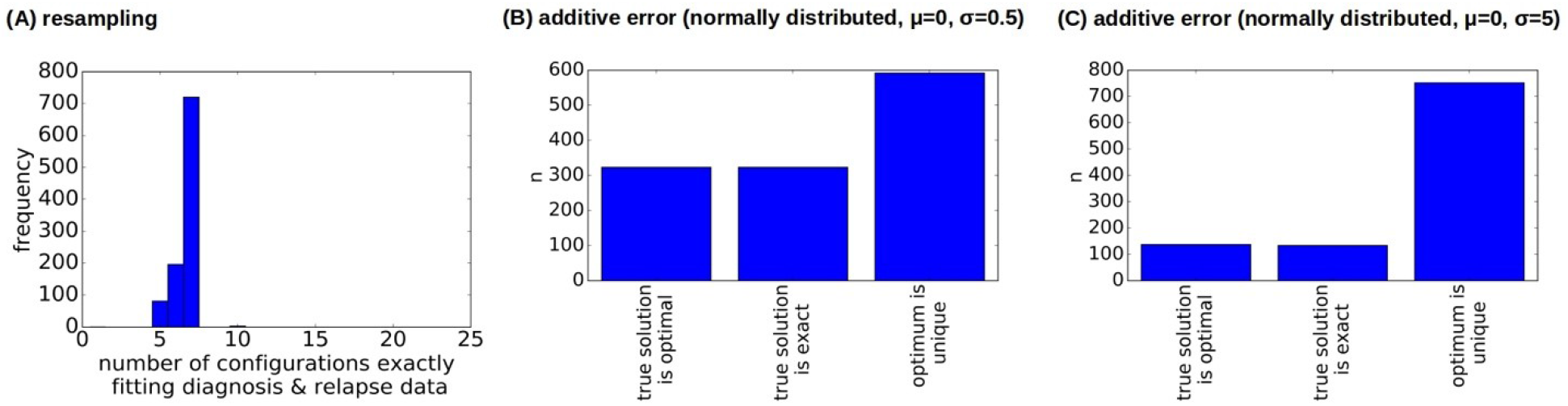
Impact of errors on the reconstructions for a sample with two founders. **(A)** Impact of sampling error on the number of clonal hierarchies compatible with diagnosis and relapse data. The vertical axis shows how many of the 1000 multinomial resamplings were compatible with 1, 2, 3, … hierarchies respectively. **(B, C)** Impact of measurement errors on the reconstructed hierarchies. We considered 1000 perturbed versions of the original data. Additive perturbations of the VAF were drawn from a normal distribution with mean zero and standard deviation 0.5 (B) or 5 (C). We observe that in the majority of cases the true configuration is not optimal.

The reason for this observation is as follows (see Fig. 9). In the exact scenario there exist two founder mutations. The frequencies of both founder mutations add up to 100%. In presence of errors it can happen that the frequencies of both founder mutations do not add up to exactly 100%. If their sum is slightly less than 100% the true hierarchy still leads to an exact solution (to compensate for the error the exact solution contains a small number of healthy cells). If due to the random error the sum over both sub-trees is slightly more than 100%, an exact solution is no longer possible. To circumvent this we can relax the dataset by artificially adding a small number of healthy cells, e.g., x% to the dataset. In this case, for measurements where the frequencies of both founding clones add up to less than 100%+x% the true configuration still is an exact solution. We see in Figure 10 that this relaxation increases the number of cases where the true solution is an optimal solution.

**Figure 9:**
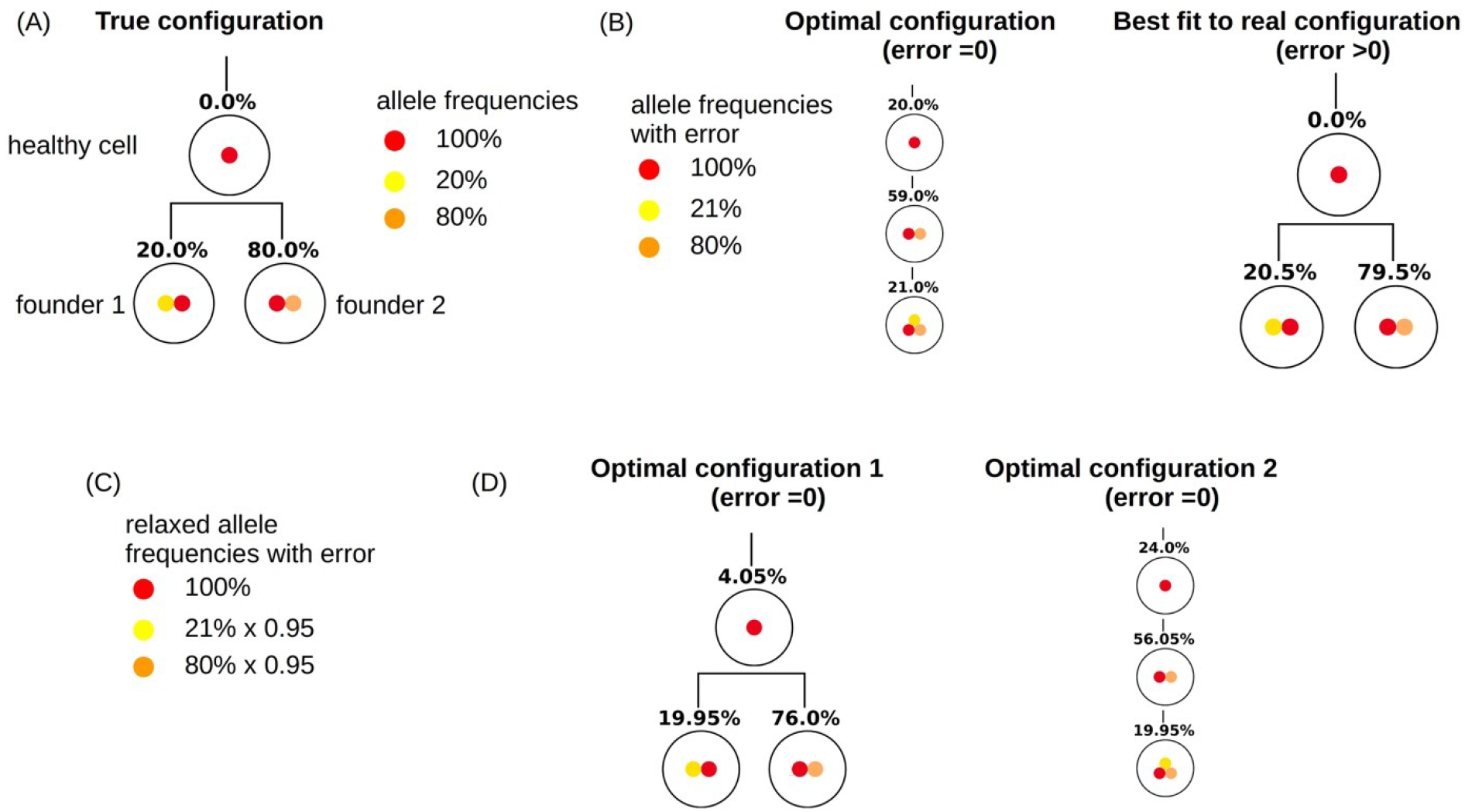
Relaxation of measurements. This Figure demonstrates how relaxation of the VAF can help to make the true configuration an optimal configuration. **(A)** True configuration with two founder clones. Wild type cells are added to the tree to have a unique root. Since the sample only contains leukemic cells the frequency of wild type cells equals zero. The frequencies of the two founder clones add up to 100%. **(B)** Variant allele frequencies are perturbed by a measurement error. The frequencies of the two founding mutations no longer add up to 100%. Therefore the true hierarchy is no longer an optimal hierarchy. **(C)** By multiplying the frequencies of mutated alleles with 0.95 we artificially add 5% of healthy cells to the sample. If we reconstruct the clonal hierarchies for the modified data the true hierarchy is among the optimal hierarchies.

**Figure 10:**
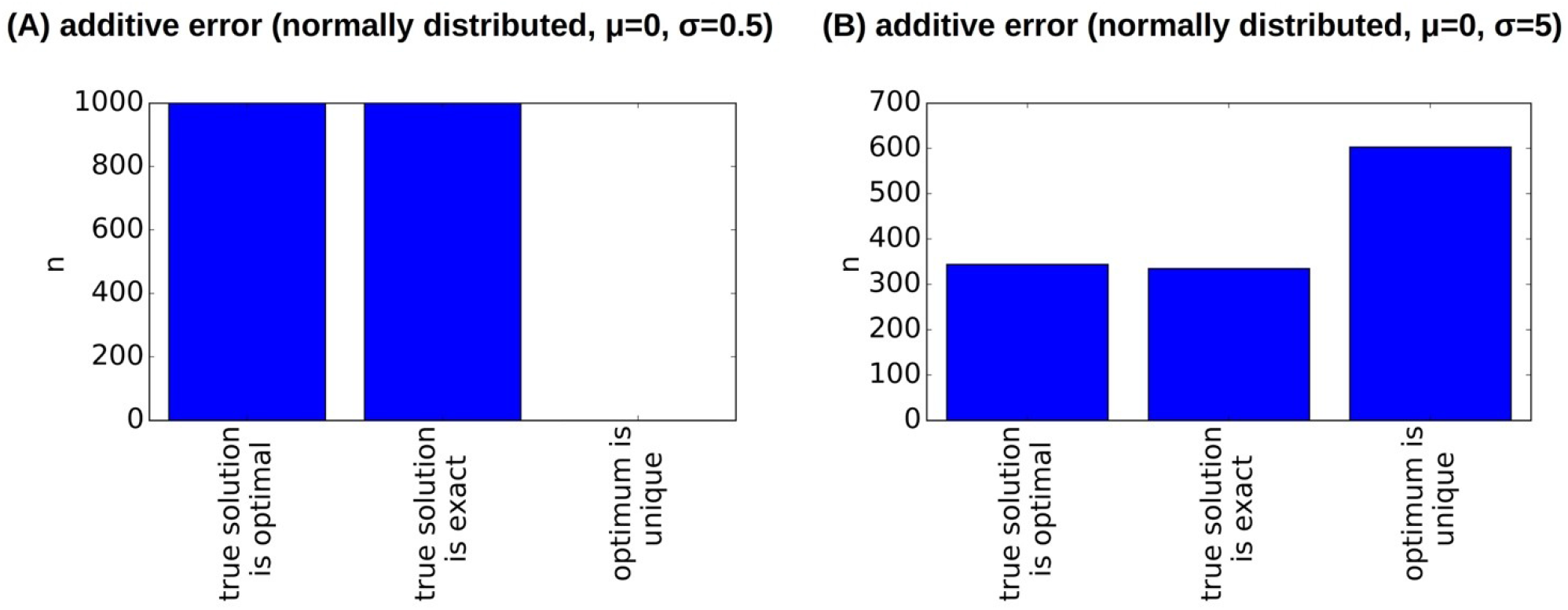
Clonal hierarchies fitting the relaxed dataset. Here we consider the relaxed version of the dataset from Patient 7. The relaxation makes the reconstruction more robust to errors. In comparison to Fig. 8 B, C the true hierarchy is in many cases an optimal hierarchy.

## Discussion

The aim of this study is to investigate the ambiguity of clonal hierarchies that are reconstructed from bulk sequencing data. For this purpose, we develop an algorithm that systematically tests which subset of all clonal hierarchies optimally fits a given dataset. We test this algorithm using bulk VAFs that have been calculated based on cell sequencing data sets. Since single cell sequencing reveals the true clonal hierarchy this approach enables us to compare the output of our algorithm to the real configuration [23, 25].

First, we assume that the input data is exact, i.e., neither sampling nor measurement errors occur. Then for most of the considered patient samples exactly one clonal hierarchy optimally fits the bulk VAF. This clonal hierarchy is identical to the hierarchy obtained from single cell sequencing. In two of the considered patients, even for exact input data more than one clonal hierarchy is compatible with the bulk allele frequencies. The true hierarchy obtained from single cell sequencing is among them. This finding implies that even in absence of measurement error, the clonal hierarchy may not be uniquely defined by the bulk VAF.

When drawing multiple samples from the same malignant cell population the variant allele frequencies may differ from one sample to another. This may be caused by sampling error, or it may reflect inhomogeneity of the tumor. Assuming the tumor to be homogeneous, we aim to quantify the impact of sampling error on the reconstructed hierarchies. For each patient, the number of sequenced leukemic single cells *n* and the frequencies *f_i_* of the different clones are known. To simulate the sampling error, for each patient we generate 1000 random samples of size *n* drawn from a multinomial distribution with probabilities *p_i_=f_i_*. For each of these random samples we calculate the bulk allele frequencies and construct all clonal hierarchies compatible with them. Based on results of this exercise we conclude that the sampling error has a negligible impact on the obtained clonal hierarchies, at least for the data at our disposal.

We test the robustness of the reconstruction by adding normally distributed errors of different amplitude to the bulk VAFs calculated from the single cell sequencing data. This takes into account potential misreads during the sequencing, amplification errors or impurities of the sample. We observe that for errors of about 5-10% the true hierarchy not necessarily remains optimal. This especially applies to data sets where the frequency of the founding clone is of the order of magnitude of the error. However, even in this case, the true clonal structure is among the upper 3-15% of the best fitting hierarchies. This implies that also in the presence of relevant errors, our algorithm allows to rule out most tree configurations and results in a small subset of possible clonal hierarchies fitting to a data sample.

Mathematical models indicate that tree characteristics, e.g., the depth of the tree, correlate with clonal properties such as self-renewal and proliferation rate [19]. In this context is can be sufficient to have an estimate of the depth of the true clonal hierarchy to draw conclusions about the effect of a mutation on cell kinetics or patient prognosis. This implies that in the case of non-unique clonal hierarchies, biological conclusions can be drawn if the potential hierarchies are sufficiently similar to each other.

Having measurements of bulk VAFs provided, our computational approach can be used to rank all possible clonal hierarchies based on their compatibility with the data (i.e., the smaller the error when fitting the dataset to a given hierarchy, the better the rank of the respective hierarchy). For all datasets considered in this study the real hierarchy is among the upper 3-15% of this ranking. Taking into account that in case of n clones n^n-1^ possible hierarchies exists our algorithm allows to rule out a significant number of them. Our algorithm can also be applied to scenarios in which the disease is derived from multiple founding clones.

Our computational approach can be used to study how sensitive the reconstructed hierarchies are to perturbations of the input data. By adding random errors to the input data obtained from an experiment and by repeating the reconstruction with the perturbed input data it turns out that some datasets are robust with respect to the perturbations. This means that the obtained optimal clonal hierarchies do not change if the input data is perturbed. For other datasets perturbations of the input data leads to a change of the reconstructed hierarchies, indicating that the reconstruction may be affected by measurement errors. The robustness of a given dataset can be checked using our proposed framework. It is straightforward to take into account that the measured frequencies may have different confidence intervals. In principle our approach can also be applied to clustered single nucleotide variants (SNVs). Since the number of detected SNVs is usually high, the variants are grouped into clusters according to their allele frequencies. Each cluster comprises all SNVs with a similar allele frequency. The cluster center is defined as the average allele frequency of all SNVs that belong to the respective cluster. In this setting our algorithm can be applied using cluster centers as input data.

Mechanistic mathematical models allow to extract relevant information from clonal hierarchies, such as estimates of proliferation rates and self-renewal of the different clones [19, 30]. Correlating these estimates with detected mutations and clinical observations may provide new insights into AML pathophysiology [9, 19]. The proposed framework is a first attempt to quantify the ambiguity emerging during reconstruction of clonal hierarchies from bulk sequencing data. It allows to identify when such reconstructions are reliable and can be used as input data for mechanistic models. This knowledge helps to make available routine clinical data to studies that require clonally resolved input [31].

## Acknowledgements

This work was supported by research funding from the German Research Foundation DFG (SFB 873; subproject B08).

## Conflict of interest statement

The authors declare that the research was conducted in the absence of any commercial or financial relationships that could be construed as a potential conflict of interest.

## Notes

https://ashpublications.org/bloodadvances/article/4/5/943/452671/Single-cell-mutational-profiling-enhances-the

### Competing Interest Statement

The authors have declared no competing interest.

